# Comparison of Three Herd-Level Surveillance Methods for Porcine Reproductive and Respiratory Syndrome Virus

**DOI:** 10.64898/2026.03.31.713744

**Authors:** Alison C. Neujahr, Todd E. Williams, Joshua L. DeMers, Bailey M. Barcal, Janel S. Peterson, Cameron S. Schmitt, Kristen K. Bernhard

## Abstract

Porcine Reproductive and Respiratory Syndrome Virus (PRRSV) represents a considerable health and economic burden for the swine production industry. highlighting the importance of disease surveillance and mitigation strategies. Accordingly, this study aimed to evaluate a novel environmental surveillance approach, referred to as DARO Systems, to detect PRRSV against established serum and oral fluids approaches. The study design consisted of one PRRSV infected seeder pig and 46 naïve nursery pigs and each surveillance approach was deployed to assess time to PRRSV detection. Findings showed the DARO Systems surveillance approach detected PRRSV 1-2 days earlier than serum and oral fluids surveillance approaches.

---

Routine pathogen surveillance testing within the swine industry serves as the key to ensuring the mitigation of biosecurity threats.^8^ As such, producers routinely test for high consequence viral and bacterial pathogens such as PRRSV. PRRSV is a highly infectious virus that causes a large economic deficit within the US swine industry of roughly 1.2 billion dollars/year due to the reduction in animal performance, high mortality rates, and treatment and prevention cost.^7,11^ Early identification of PRRSV within a facility can result in efforts to mitigate spread, such as vaccination and closure of barns.^16^

There are numerous ways to conduct surveillance for PRRSV in swine herds including partial herd sampling (i.e. the collection of samples from a subset of animals or pens to infer population-level infection status) by testing serum, oral fluids, processing fluids, and tongue tips. However, there are numerous pitfalls to current PRRSV surveillance strategies, including that some sampling approaches are not applicable across all life stages (processing fluids) or they are difficult to scale and do not sample the entire herd (serum, tongue tips, and oral fluids). Serum testing and oral fluids approaches are the most widely used approaches as they span life stages and do not depend on deceased animals. Venepuncture for serum collection from swine is tedious and stressful for the hogs and samplers. Additionally, scalability is difficult as labour and cost limitations limit feasibility in large operation settings as the number of hogs that would need to be bled to detect PRRSV when the prevalence is low is substantial.^14^ Therefore other surveillance techniques that test larger proportions of the herd, including oral fluids, are used.^3,10^ Oral fluids is a simple, non-invasive sample collection method that allows pigs to chew on an undyed cotton rope and saliva is subsequently tested for the presence of pathogens.^5^ Oral fluids can be collected on an individual or pen-based level.^9^ While oral fluid collection mitigates labor challenges and animal welfare concerns, it introduces a potential sampling bias toward healthy sub-populations. As a result of the method being reliant on active engagement with the chew rope, lethargic or morbid animals often fail to contribute a sample.^6^ This behavioral discrepancy likely leads to an under-representation of PRRSV infected swine, thereby underestimating prevalence estimates and increasing the risk of false-negative reporting.^13^

To mitigate the consequences of PRRSV infection in herds, disease surveillance needs to be fast, reliable, simple, and needs to sample all animals. This study reports a novel testing procedure, known as DARO, a non-invasive herd-level fecal collection that can be conducted by any skill-level farm employee and is coupled with a RT-qPCR workflow designed to improve detection of viral RNA from heterogenous environmental matrices. The purpose of this study was to evaluate the differences in PRRSV detection using three different herd-level, age-agnostic surveillance methods: oral fluids, serum, and DARO.

## Animal care and use

This study involved non-invasive environmental fecal sampling where review and approval by the DARO Institutional Animal Care and Use Committee (IACUC) was determined to be not required. Serum and oral fluid sampling was reviewed and approved by Pipestone Research IACUC #2025-14 and the study was conducted according to the Guide for the Care and Use of Agricultural Animals in Research and Teaching.^4^

## Materials and Methods

### Study Design

The study was conducted within an Animal Biosafety Level (BSL 2) barn facility, with six independent rooms and air spaces and a metal tri-deck flooring over a 4’ cement pit located in Minnesota. One room within the facility was used. The study consisted of 46 PRRSV naïve nursery pigs housed in 6 Pens. On day 0 of the trial, one seeder pig, was inoculated with PRRSV wild type strain (1.4.4 L1C) prior to being placed in Pen 1. The virus originated from a sow farm with an active PRRSV outbreak in Minnesota (2021). South Dakota University Animal Disease Research and Diagnostic Laboratory propagated the virus where the viral concentration was 1×103.5 TCID/50mL. The pig was given 2mL administration of the propagated virus via intermuscular injection. No pigs from Pen 1 were sampled to ensure there was no sample collection from the fecal excretion and oral fluid of the seeder pig. Pens 2 through Pen 6 were surveyed for the 7-day duration of the study. A total of 39 pigs were housed in Pens 2-6 with the following per pen values: Pen 2-8 pigs, Pen 3-8 pigs, Pen 4-8 pigs, Pen 5-8 pigs, Pen 6-7 pigs. Hogs were monitored using three different surveillance methods (oral fluid, serum, and DARO Systems) to observe infection spread and compare time to detection of the different surveillance approaches. All surveillance methods were utilized daily for the duration of the 7-day study.

### Sample Collection and Processing

Daily oral fluid samples were collected from designated pens (between Pen 2 & 3 and Pen 4 & 5) representing 82% (33/39 animals) of the surveyed herd having access to the rope (excluding Pen 1 and Pen 6) using the swine producers herd management protocol (Figure 1). Rope placement was selected to ensure there was no collection from the seeder pig (housed in Pen 1) and Pen 6 was excluded from oral fluid testing to mimic operating barns standard practice of providing oral fluid rope to only a portion of all animals. Animals engaged with the hung rope for 20 minutes. Both ropes were individually squeegeed to remove the oral fluid and collected into a single 5 mL Falcon tube (Corning, NY, USA), respectively. Serum was collected daily according to routine swine producers herd management protocol. Serum was drawn at random from two pigs in Pen 2 and Pen 4, and from three pigs in Pen 3 and Pen 5, sampling 26% (10/39 animals) of the surveyed herd. A total of 2 pig serum samples were pooled daily from Pen 2 and Pen 4, and a total of 3 pig serum samples were pooled daily from Pen 3 and Pen 5. To replicate routine serum surveillance strategies, serum was pooled by pens (Pens 2 & 3 combined and Pens 4 & 5 combined) (Figure 1) prior to sample processing.

**Figure 1.**
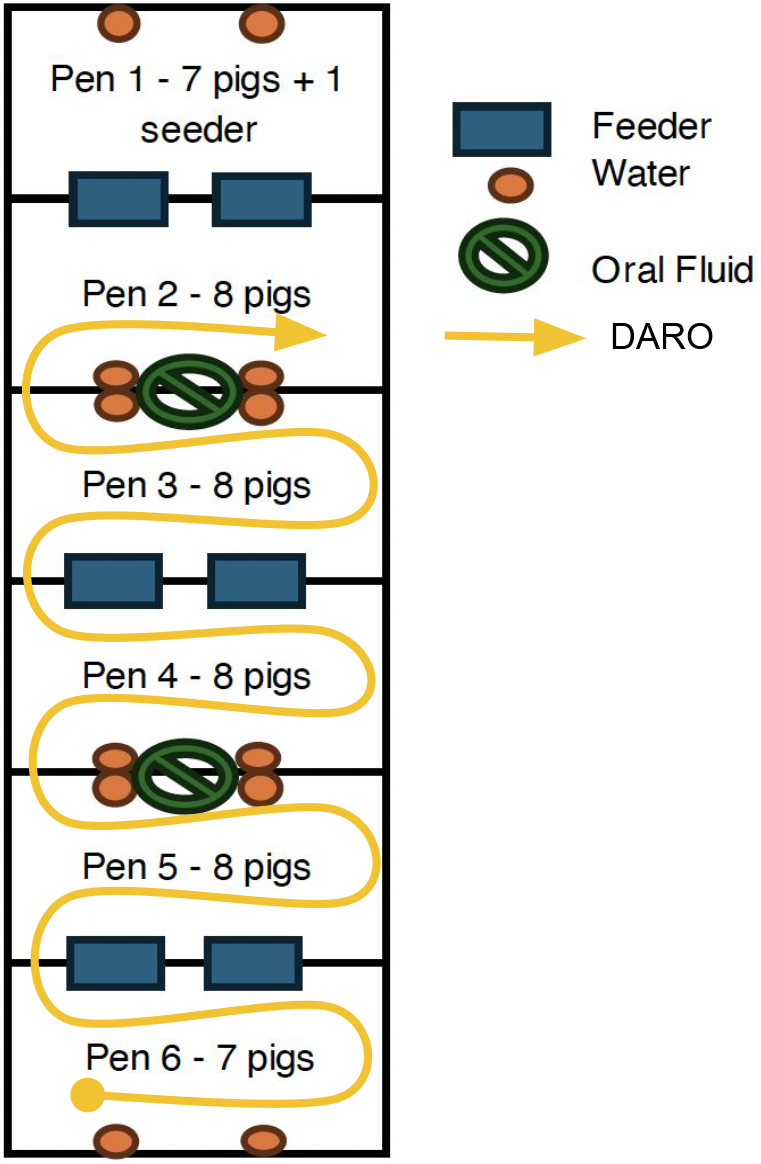
Schematic layout of study design.

Oral fluid and serum samples were sent to Iowa State University Vet Diagnostic Lab (ISU VDL) and tested for the presence of PRRSV using the same RT-qPCR approach according to the standard ISU VDL protocol. Samples were considered positive or negative by the lab reported cutoff of a Ct value of ≤ 37.

Daily fecal samples were collected using DARO collection methodology (Pending Patent #19/256,939). Trained collection staff personnel obtained an aggregated near-whole-herd fecal sample by comprehensively walking all pens and dunging areas with absorbent shoe covers starting at Pen 6 and moving through 50% of Pen 2 (the side closest to Pen 3) until all pens (except Pen 1 with the seeder pig) had been collected (Figure 1). Partial collection of Pen 2 and no collection of Pen 1 was conducted to reduce collection of excretion from the seeder pig within Pen 1. All 5 collected pens were aggregated into one sample. Samples were shipped with a cold pack to DARO Laboratory for processing for the presence of PRRSV using patent pending DARO System’s sample processing technique (Pending Patent #19/256,939). Nucleic acid was extracted from the samples using the MagMAX Wastewater Ultra Nucleic Acid Kit (Applied Biosystems, Foster City, CA, USA) according to manufacturer protocol. Xeno (VetMAX Xeno internal positive control, Applied Biosystems, Foster City, CA, USA) was added as an internal positive control to each sample at a concentration of 20,000 copies per reaction post lysis. Extracted samples were prepped for quantification using the VetMAX PRRSV 3.0 Reagent Kit (Applied Biosystems, Foster City, CA, USA) following the manufacturers protocol. Each RT-qPCR run included an extraction and template negative control and positive control. All samples in the study were considered positive with a Ct value of ≤ 37.

## Results

The objective of this descriptive study was to determine the time to detection of PRRSV using three different surveillance collection methods PRRSV was first detected with the DARO approach on day 3 while PRRSV was detected in serum on day 4 and oral fluids on day 5 (Table 1). All sampling methods continually detected PRRSV with decreasing CT values (increasing viral load) following the first of detection, consistent with progressive PRRSV infection in pigs(Figure 2, Table 1).

**Table 1.**
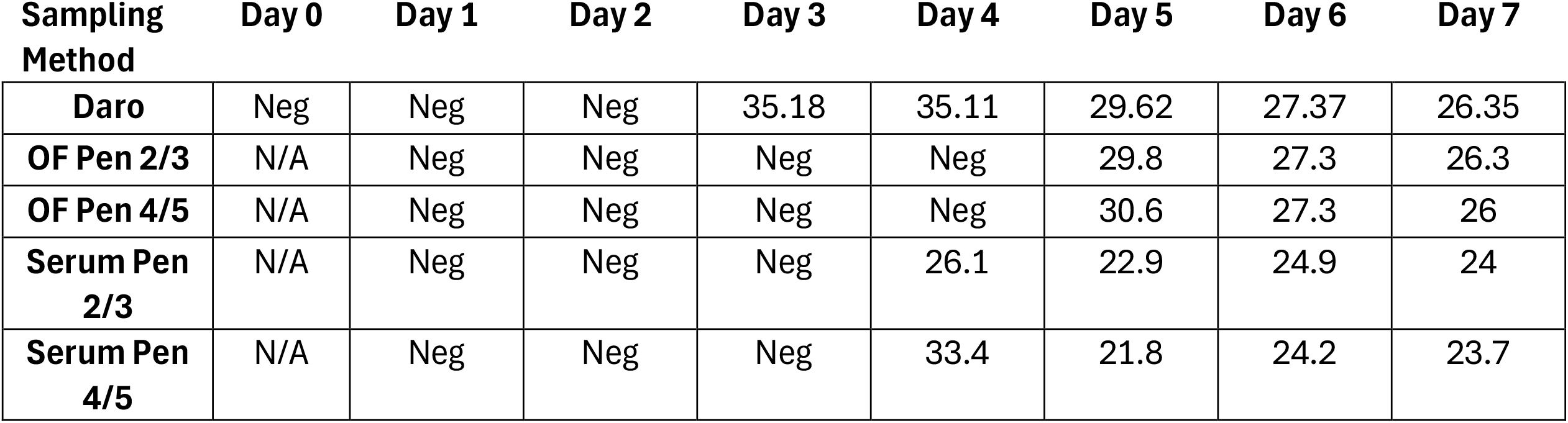
CT value results for each testing method used in the study. NEG= PRRSV not detected by RT-qPCR. N/A= not sampled NEG= PRRSV not detected by RT-qPCR. N/A= not sampled.

**Figure 2.**
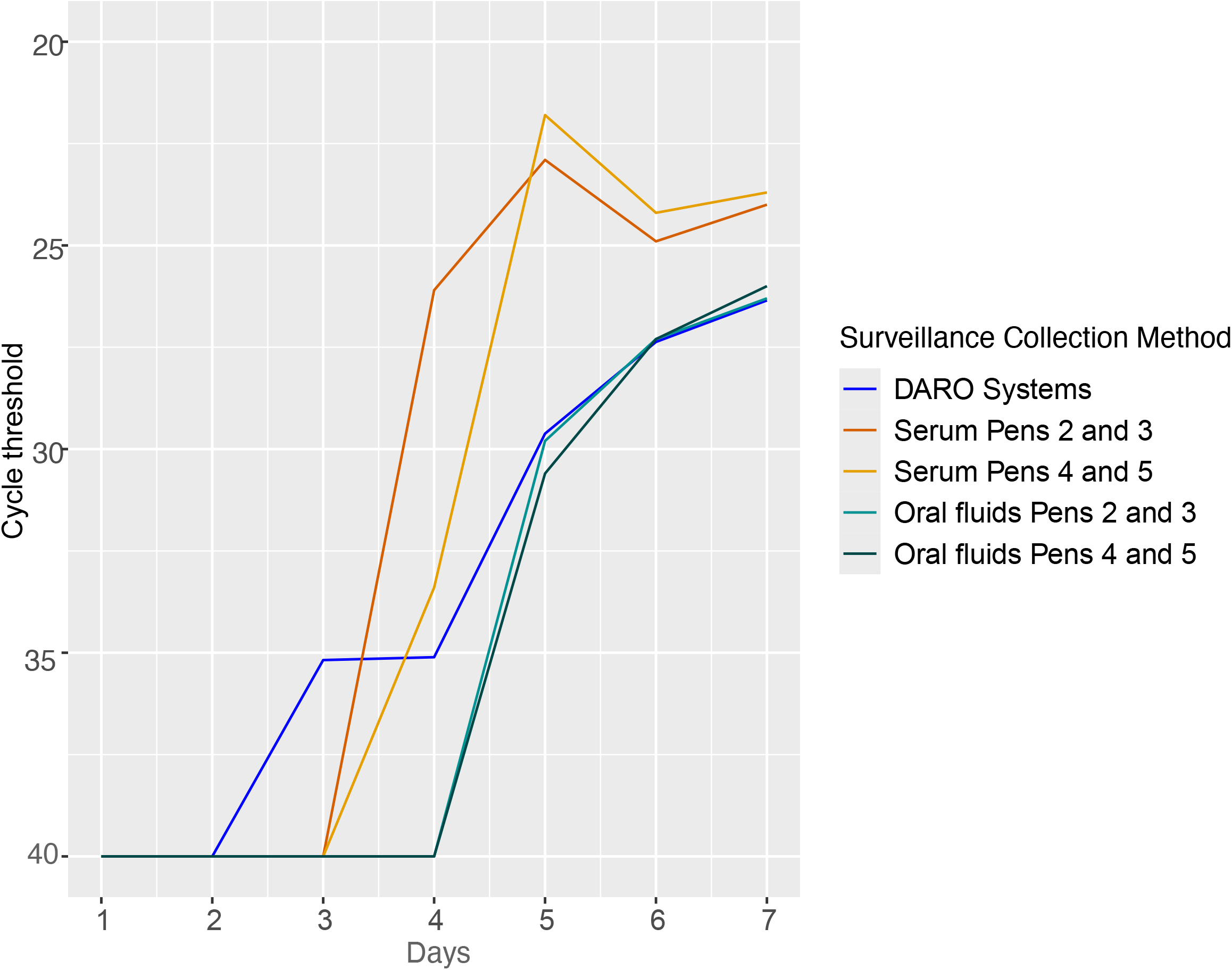
Days to positive PPRSV detection comparing DARO Systems, serum, and oral fluid sampling. DARO Systems identified PRRSV one day earlier than serum and two days earlier than oral fluids.

## Discussion

Comprehensive herd-level collection methods, such as DARO aggregated fecal sampling method, offers advantages over partial herd sampling approaches (e.g., oral fluid or serum) for detecting shedding individuals. Such practices allow for the minimization of risk of not detecting a single infected animal due to chance or sampling bias. As a result, comprehensive herd-level collection ensures earlier detection of infected individuals within a given herd. Moreover, reports show population testing provides additional advantages such as significant per capita labor and cost savings at scale and allows for testing in large populations where individual testing capabilities are limited.^12^ Similarly, DARO leverages a comprehensive herd-level collection method to offer early-detection when individual testing is cost and labor prohibitive. In a controlled observational study of a small herd (n=39 sampled animals), DARO identified PRRSV one day earlier than serum and two days earlier than oral fluids.

More comprehensive sampling with the DARO collection approach likely led to early detection of PRRSV in this study, as opposed to the detection method being more sensitive from environmental sampling opposed to serum or oral fluids. Two observations lead to this conclusion: 1) using environmental fecal sampling, nearly 100% of the herd was sampled, where 82% of the surveyed herd had access to a rope for oral fluid collection and 26% of the surveyed herd was bled daily, and 2) CT values continuously decreased for each sampling methodology and CT values were lower in serum samples compared to both DARO and OF, indicating the sensitivity of the assays did not explain detection differences. Serum samples having lower CT values are expected given that pools were created from either 2 or 3 serum samples, whereas testing from DARO and oral fluids included samples from dozens of animals in each pool.

Sampling limitations of current herd-wide surveillance strategies are exacerbated when applying surveillance approaches to real-world herd operations (e.g., commercial operations housing ∼1,000 pigs within a room). Frequent sampling of individual pigs becomes cost, labor, and pig-behavior prohibitive. The efficacy of DARO to detect PRRSV within a large commercial setting was demonstrated where DARO identified PRRSV on average 3.91 days earlier than oral fluid testing in nursery barns consisting on average of ∼1,560 head.^1^ The reduced capability of partial herd sampling (e.g., oral fluids, serum) is particularly evident in emerging disease situations where herd prevalence is low and therefore difficult to detect. Introduction of PRRSV likely occurs from a single pig, thus the likelihood of detection during early transmission phases increases as the proportion of pigs sampled increases. As DARO tests environmental samples from an entire barn as opposed to samples collected from pigs themselves, the likelihood of early detection improves. The early detection found from DARO collection method identified PRRSV infections sooner during an outbreak scenario, thus more accurately reflecting the health status of the entire herd. A shift towards comprehensive herd-level collection approaches can improve PRRSV surveillance in US swine production.

### Limitations

The major limitation of the study is that it is a single-replicate study design, and therefore formal inferential statistical methods cannot be applied to the results. Although the results are descriptive, the conclusion DARO surveillance strategy was able to detect PRRSV infection in the herd earlier than serum and oral fluids due to increased sampling fraction is well-reasoned. This study allowed for a controlled environment to observe PRRSV spread and compare and contrast surveillance method capabilities at the known start of an emerging PRRSV outbreak and the results justify a larger comparative study to more thoroughly assess differences between approaches.

## Future Directions

Based on the results of this study, a larger study comparing herd-wide surveillance strategies in larger barns that more accurately reflect sampling approaches of producers is needed. Additionally, developing a surveillance system that can be deployed to all stages of pig production across all barn configurations is crucial to ensuring every stage of swine production can reliably be surveyed for pathogens. All production stages (breeding, nursery, and grow-finish) require PRRSV surveillance because infection occurs across age groups, causing reproductive disease in sows and respiratory disease in growing pigs. In addition, epidemiological modeling shows that PRRSV transmission occurs across interconnected production phases—including breeding, nursery, and finisher systems—through animal movement, local transmission, and reinfection dynamics. Reports show that sows kept in groups are less interested than grow-finish pigs and collection from young piglets is not feasible as training rate for rope chewing is near 0%.^15^ Thus, a clear future direction of this work is to determine the feasibility of DARO to be used across different life stages. Additionally, the DARO approach is applicable to most relevant pathogens and future studies will expand the repertoire of pathogens that can be tested for.

Implications Under the conditions of this study:

- Representation of barn health is shown with comprehensive herd-level testing.
- Earlier detection window allows for mitigation planning.
- DARO Systems allows for scalability.

## Acknowledgments

The authors thank Dr. Joseph Fauver for his helpful edits and comments to the manuscript.

## Disclaimer

Scientific manuscripts published in the *Journal of Swine Health and Production* are peer reviewed. However, information on medications, feed, and management techniques may be specific to the research or commercial situation presented in the manuscript. It is the responsibility of the reader to use information responsibly and in accordance with the rules and regulations governing research or the practice of veterinary medicine in their country or region.

